# Misinterpreting the horseshoe effect in neuroscience

**DOI:** 10.1101/2022.03.04.482986

**Authors:** Timothée Proix, Matthew G. Perich, Tomislav Milekovic

## Abstract

Dimensionality reduction methods are frequently used to analyze high-dimensional activity of cortical neuron populations during behavior. The resulting oscillatory trajectories that consistently emerge from this analysis have been interpreted as a signature of latent dynamical systems. Here, we show that these oscillatory trajectories necessarily result from applying dimensionality reduction methods on recordings that approximately exhibit continuous variation in time, regardless of whether or not the recorded system incorporates latent dynamics.

## Main text

Modern experimental techniques allow researchers to monitor the activity of hundreds to thousands of neurons simultaneously. Analyzing, interpreting and understanding how such high-dimensional neural activity evolves over time is challenging. Dimensionality reduction methods are commonly used to display this activity by representing it as trajectories within a lower-dimensional subspace^1,2^. Yet, these methods consistently produce oscillatory trajectories when applied to the activity of neural populations in many experimental scenarios, including the motor cortex during movement^3–5^, prefrontal cortex during working memory^6^, and lateral sensorimotor cortex during speech articulation^7^. The shape of these trajectories can be stable across recorded neuron populations, tasks, trials, and time^8^. Due to their consistency, these oscillatory low-dimensional neural trajectories have been interpreted as evidence that the activity of cortical populations behaves as a latent dynamical system. Here, latent dynamical system means that the evolution of the majority of the neural population activity can be described by a differential equation of the past neural activity:

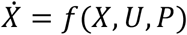

with X the activity of all neurons concatenated into a vector, P behavioral parameters, and U external inputs. This is in contrast with the hypothesis that the majority of the neural activity can be described solely as a function of behavioral parameters and input stimuli^3,9–11^:

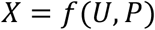

Oscillatory trajectories also emerge when applying dimensionality reduction methods to high-dimensional data in other fields, such as genetics^12^, ecology^13^, climatology^14^ and time series analysis^15^. In contrast to seeking deeper insight, these fields recognize oscillatory trajectories as a product of the “horseshoe effect”, a mathematical rule stating that oscillatory trajectories necessarily emerge when applying dimensionality reduction to data exhibiting continuous variation in time or space. This rule is valid for numerous commonly-used dimensionality reduction methods, including principal component analysis (PCA)^10,11,16^, multidimensional scaling^17^, isomap^17^, local linear embedding^17^, and kernel PCA^17^.

To appreciate the horseshoe effect, consider a Society for Neuroscience meeting. Thousands of neuroscientists throughout the world have gathered for a poster session in a hallway. The hallway is divided into different sectors corresponding to different topics. While crossing the poster session’s hallway at a constant speed, we record neuroscientists’ nationality. We observe that neuroscientists from the same country tend to similarly group together around the same topic. For six countries, we compute the number of neuroscientists as a function of time during our crossing of the hallway and apply PCA (Fig. 1a). Regardless of the spatial ordering of data, the PCA finds the orthogonal linear combinations of nationality records, called “modes”, that explain the variance in decreasing order. The first mode explains the most variance of the whole dataset, the second mode the most variance of the full dataset minus the first mode, etc. Intriguingly, the time evolution of the modes, called “latent variables”, is oscillatory (Fig. 1b).

**Figure 1.**
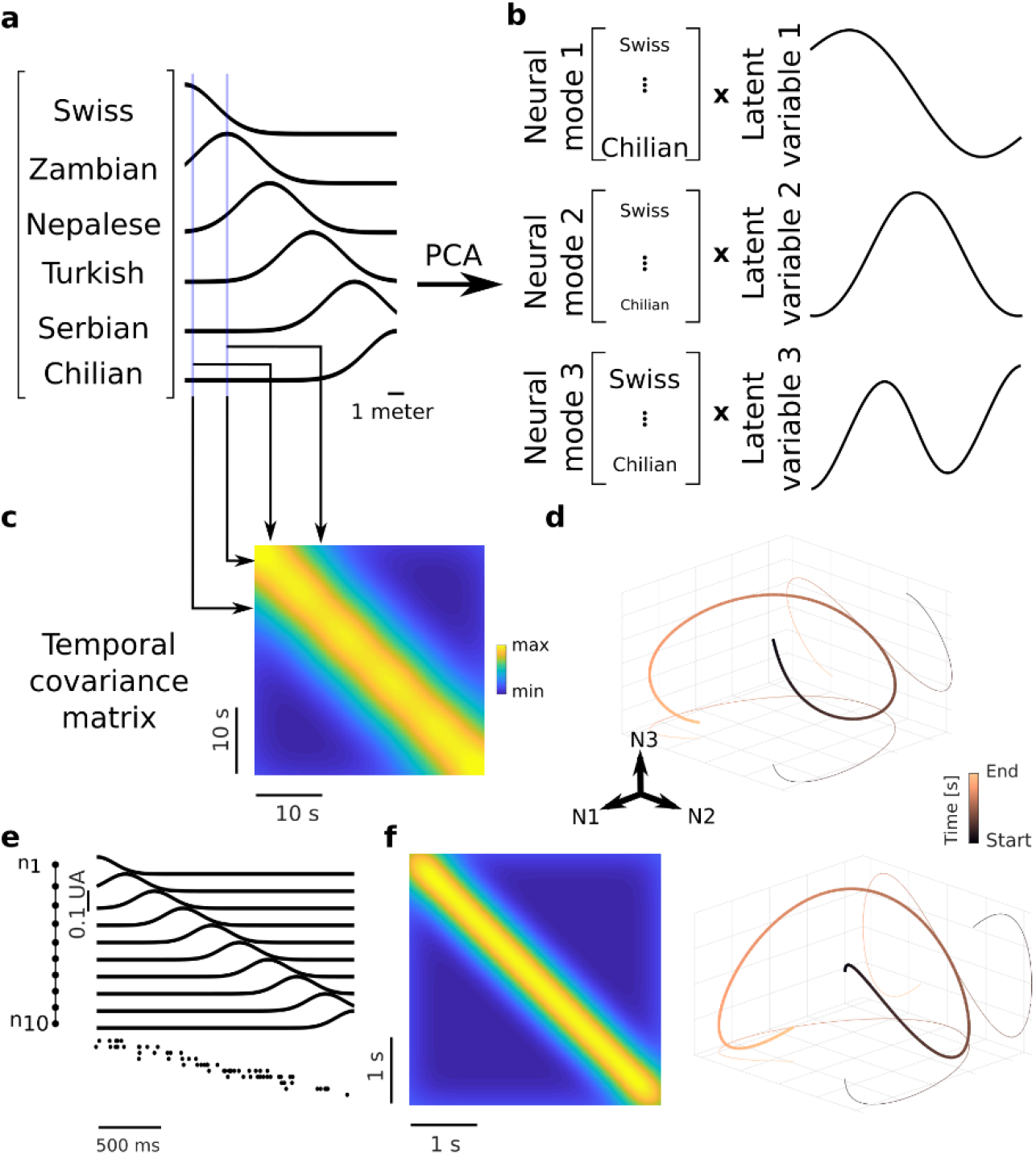
Latent variables of dimensionality reduction methods necessarily trace oscillatory trajectories if the data exhibit continuous variation in time regardless of the dynamics underlying the data. (**a**) Plots show number of neuroscientists from six different nationalities as a function of the time of the experimenter crossing the hallway. (**b**) PCA applied on this dataset derives neural modes and latent variables. Due to the horseshoe effect, latent variables approximate the Fourier series (sinusoidal curves of increasing frequencies). (**c**) Computing the covariance between all pairs of vectors containing the number of neuroscientists at one instant gives the temporal covariance matrix, which for our dataset has a Toeplitz structure. (**d**) The first three latent variables, which are approximately the first three Fourier series components, trace the signature horseshoe curve. (**e**) Simulated activity using the line neural population model with 100 neurons. Only ten neurons are shown for visualization. (**f**) Temporal covariance matrix is Toeplitz (left panel) and, due to the horseshoe effect, the three-dimensional neural trajectory traces the horseshoe curve (right panel).

Ultimately, for this dataset, the latent variables will necessarily approximate the Fourier series: cosines and sines ordered by multiples of base frequency, which is the inverse of the recording duration. The logic is as follows. In our data, the “set” – the collection of unordered elements – of all the pairwise absolute differences between the number of neuroscientists for a nationality, *n*_*i*_*(t)*, collected at two given experimental time instants t_1_ and t_2_, S(*t*_*1*_, *t*_*2*_)={|*n*_*i*_*(t*_*2*_) – *n*_*i*_*(t*_*1*_)| for all nationalities *i*}, only depends on the experimental time difference τ=|t_2_ – t_1_|, and not on the particular values of t_1_ and t_2_. That is, how neuroscientist of the same nationality group together in the hallway does not depend on their nationality. This feature of the dataset is called “continuous variation in time”. It is best visualized using a temporal covariance matrix that displays the covariance of two sets of nationality measurements for all experimental time pairs (t_i_, t_j_). Because of the continuous variation in time, the temporal covariance matrix has a particular structure known as Toeplitz matrix^18^ (Fig. 1c). It can be shown analytically (Supplementary text 1) that the eigenvectors of the Toeplitz matrix are given by approximate Fourier series^18^. Since the eigenvectors of the temporal covariance matrix are proportional to the PCA latent variables, a dataset with a Toeplitz temporal covariance matrix necessarily has approximate Fourier series as its latent variables. Other dimensionality reduction methods, including multidimensional scaling, isomap, local linear embedding, and kernel PCA, have a similar relationship between the temporal covariance matrix and their latent variables, thus resulting with approximate Fourier series latent variables when the temporal covariance matrix is Toeplitz^17^.

Consequently, the latent variables of our nationality dataset are approximate sinusoidal curves of increasing frequency. Latent variables of the first three modes trace the signature horseshoe curve^16^ (Fig. 1d). This curve closely resembles oscillatory trajectories often obtained when applying dimensionality reduction methods to activity of neural populations^3,4,19^. Yet, these oscillatory trajectories are the consequence of neuroscientists being regularly tuned to locations of their nationality-preferred topic as we walk across the hallway. The position of each neuroscientist does not depend on position of other neuroscientists. The only required condition for oscillatory trajectories is the continuous variation in time of the dataset, which does not inform us about the dynamics of the underlying system generating it.

Another way to illustrate the emergence of oscillatory trajectories though the horseshoe effect is by simulating neural spiking activity. To this end, we simulated 100 neurons arranged on a line. Firing rate of each neuron is tuned to a time instant proportional to the location of the neuron on the line (Fig. 1d). Note that this dataset can be generated with two physical implementations: either by setting delayed connections between each neuron and a time source (no dynamical systems involved), or by considering a propagating wave of neural activity along the neurons’ line (represented by a dynamical system). With this model, we obtain a Toeplitz temporal covariance matrix and reproduce the approximate Fourier series of latent variables as predicted by the analytical derivation of the horseshoe effect. As in the model, the three-dimensional neural trajectory traces the horseshoe curve (Fig. 1e).

Due to noise, random or biased sampling, real-life neural datasets will never exhibit ideal continuous variation in time even if the population exhibits this property. Therefore, the temporal covariance matrix will deviate from the Toeplitz structure. Nonetheless, the similarity of the three-dimensional neural trajectories to the horseshoe curve is proportional to the similarity of the temporal covariance matrix to the Toeplitz matrix. To show this, we quantified the similarity of the temporal covariance matrix to the Toeplitz matrix using Pearson’s correlation measure R^2^; and the similarity of the four-dimensional neural trajectory to the four-dimensional horseshoe curve using canonical correlation (CC)^20^. We then simulated neural population activity recordings with different levels of noise, or using sparse random or biased sampling. R^2^ and CC remained proportional across different levels of white noise (Extended Data Fig. 1). Both R^2^ and CC values remained high for random and biased sampling, even for as few as ten randomly sampled neurons (Extended Data Fig. 2, 3). These simulations show that oscillatory trajectories will emerge from the dimensionality reduction analysis even if the data recordings are noisy, sparse or biased.

Microelectrode array recordings of the neural firing rates in the cortex approximately exhibit continuous variation in time. Therefore, due to the horseshoe effect, recorded neural activity projected into leading modes traces oscillatory trajectories that approximate the Fourier series^10,11,21–26^. To show this, we considered several multielectrode array datasets recorded from cortices of non-human primates who performed various visual or motor tasks. We first analyzed the spiking activity of a rhesus macaque monkey implanted with two 96-channel microelectrode arrays in the primary motor (M1; Fig. 2a-b) and premotor dorsal cortex (PMd; Fig. 2e-f)^27^, and another monkey implanted with a 96-channel microelectrode arrays in the primary somatosensory cortex (S1; Fig. 2i-j)^28,29^. The monkeys performed an instructed-delay center-out task using a manipulandum, with eight reaching directions. The trial-averaged temporal covariance matrix for a single reaching direction resembled a Toeplitz matrix (Fig. 2c,g,k; R^2^=0.74, 0.65 and 0.66 for M1, PMd and S1). Due to the horseshoe effect, three leading latent variables approximated the Fourier series and traced oscillatory trajectories that resembled horseshoe curves (Fig. 2d,h,l; CC=0.92, 0.92 and 0.83 for M1, PMd and S1). We then analyzed the spiking activity of a rhesus macaque monkey implanted with a microelectrode array in the V4 area of the visual cortex^30,31^ (Fig. 2m-n). The monkey performed a fixation task while a sequence of static gratings and plaid stimuli were shown peripherally in a location centered on the V4 receptive fields. Each fixation included 10 flashed stimuli shown in a one second sequence for 100 ms each. The temporal covariance was similar to a Toeplitz matrix (Fig. 2o; R^2^=0.55) and, due to the horseshoe effect, three leading latent variables also traced oscillatory trajectories that resembled the horseshoe curve (Fig. 2p; CC=0.80). Since these oscillatory trajectories are a direct consequence of the continuous variation in time of the dataset, oscillatory trajectories cannot be used as evidence that the neural activity was generated by latent dynamical systems.

**Figure 2.**
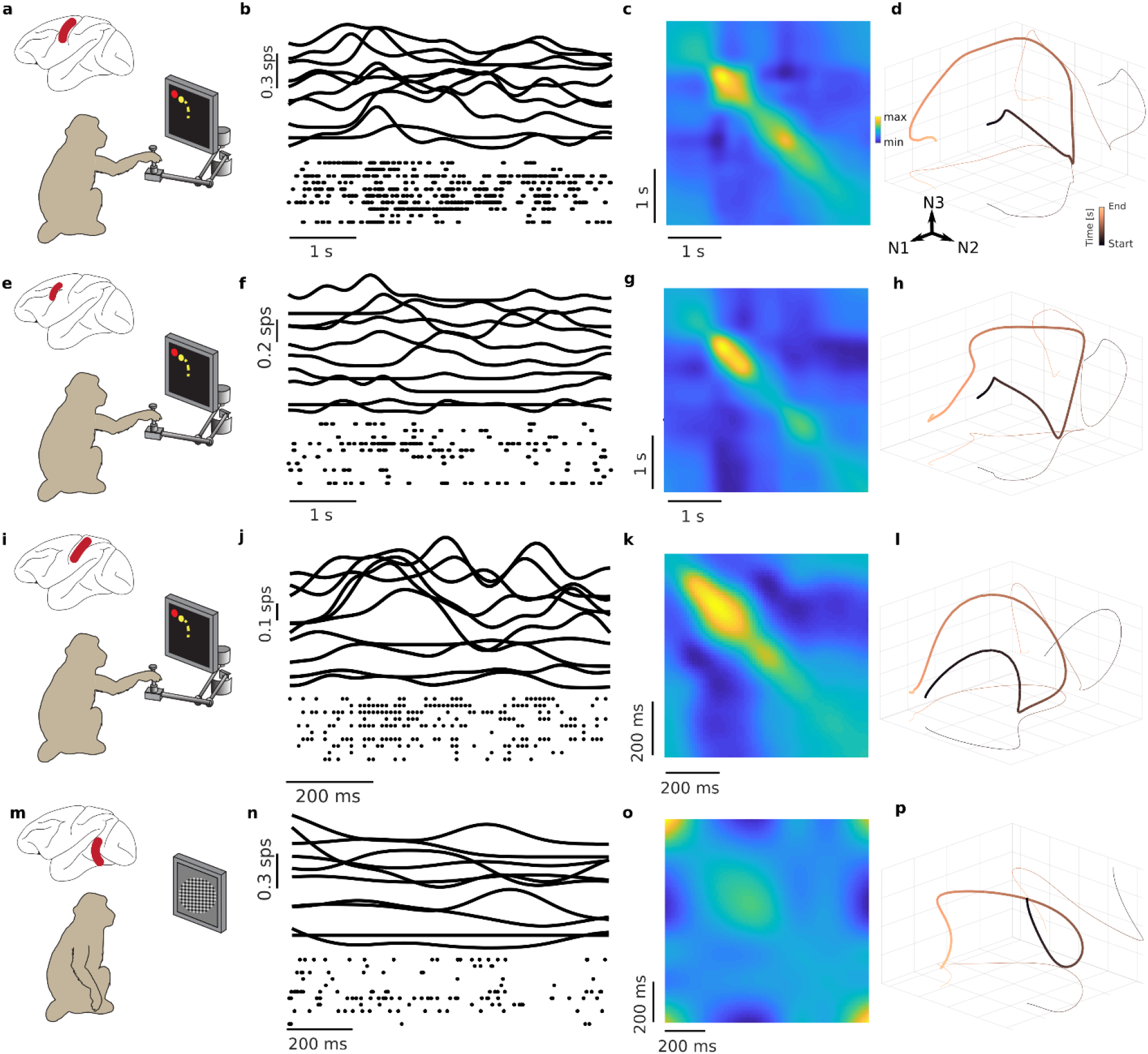
Due to the prevalence of continuous variation in time in spiking activity of cortical neural populations, dimensionality reduction techniques yield latent variables that trace oscillatory trajectories. (**a**) Spiking activity was recorded from the motor cortex of a monkey performing a reaching task. (**b**) Top panel shows the firing activity for 10 example neurons, and bottom panel shows the corresponding spiking activity. (**c**) Temporal covariance matrix of all 84 neurons approximates a Toeplitz matrix. (**d**) Due to the horseshoe effect, the three leading latent variables are approximately the first three Fourier series components. Therefore, neural activity projected in a space spanned by the three leading modes (N1, N2 and N3) traces oscillatory trajectories. (**e-h**) Similar plots to (a-d) for the same monkey and the same task showing recordings from the premotor cortex instead. (**i-l**) Similar plots to (a-d) for another monkey and the same task showing recordings from the primary somatosensory cortex. (**m-p**) Similar plots to (a-d) recorded from the visual cortex area V4 of a monkey performing a fixation task with visual stimuli. Monkey images adapted from SciDraw.io.

In datasets considering multiple tasks or conditions, sequence of neural firing will vary across tasks/conditions even if the neural activity recordings exhibit continuous variation in time. The temporal covariance matrix then has a block structure. The diagonal blocks describe the intra-condition temporal covariances and are well approximated by the Toeplitz matrix. The out-of-diagonal blocks describe the inter-condition temporal covariances. These blocks do not have a Toeplitz structure. Applying the dimensionality reduction techniques to this data results with a collection of oscillatory trajectories. This is evidenced by applying PCA on our M1 center-out dataset containing two opposite reaching conditions. The three-dimensional neural trajectories traced two near-orthogonal oscillatory trajectories, reflecting the structure of the inter-condition blocks (Fig. 3a). Applying PCA on the full M1 center-out dataset containing 8 reaching conditions resulted with three-dimensional neural trajectories tracing eight oscillatory trajectories arranged by the reaching angle (Fig. 3b). Each block of the temporal covariance matrix corresponds to a pair of reach direction, and blocks of neighboring directions are more similar than blocks of distant directions. Our line neural population model cannot reproduce these features. Nonetheless, this structure can still be replicated by a simple extension of this model that doesn’t rely on latent dynamics.

**Fig. 3.**
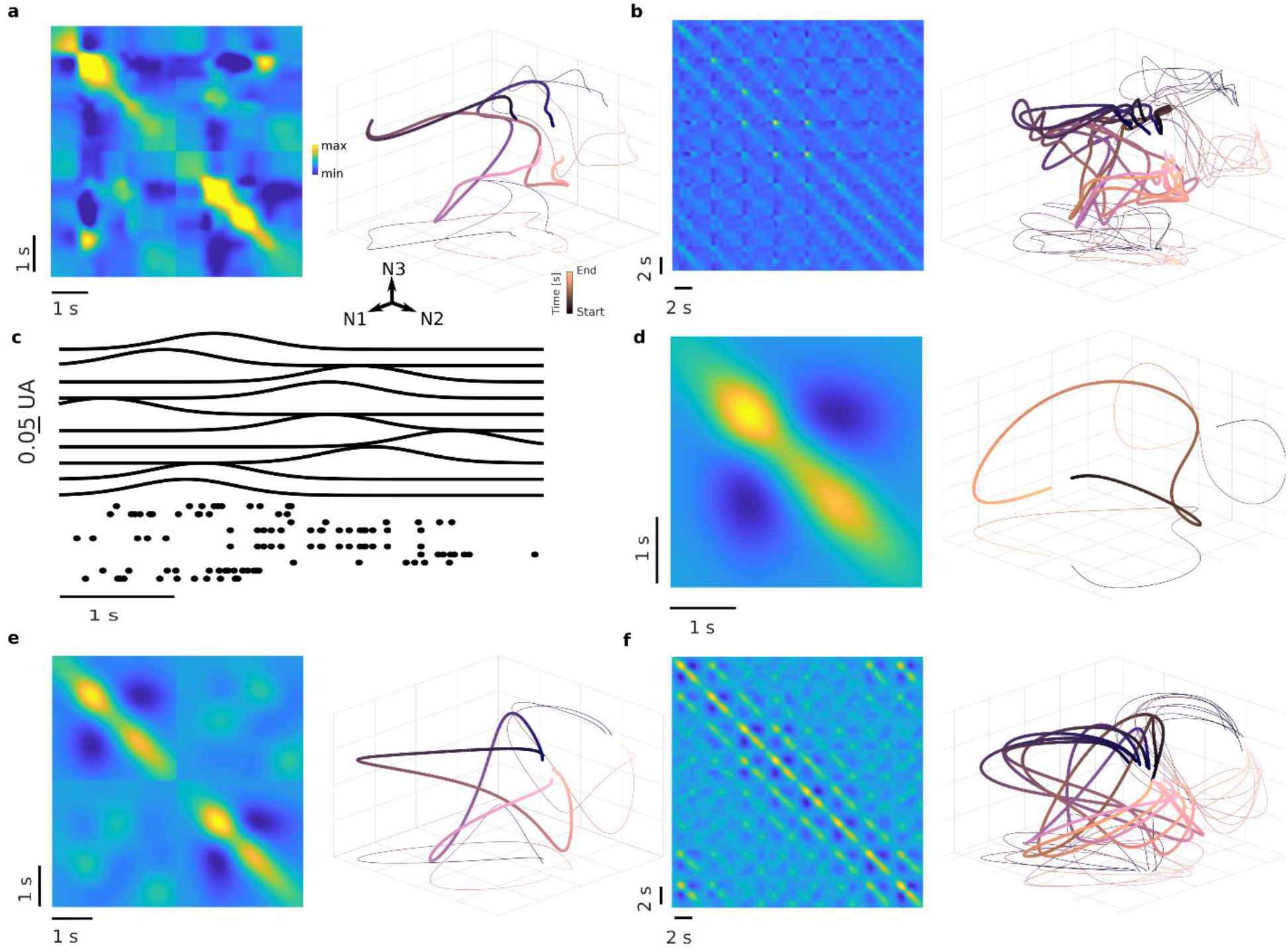
Neural population datasets containing multiple tasks and exhibiting a collection of oscillatory trajectories can be simulated by a neural population model without latent dynamics. (**a**) Applying PCA on the M1 dataset with two opposite reaching directions results with two near-orthogonal oscillatory trajectories. Colorplot shows the temporal covariance matrix of the dataset. Three-dimensional plots show the neural trajectories of the two reaches projected in the space spanned by three leading modes. (**b**) Applying PCA on the M1 dataset with all eight reaching directions results with eight oscillatory trajectories arranged by the reaching angle. Colorplot shows the temporal covariance matrix of the dataset. Three-dimensional plots show the neural trajectories of all eight reaches projected in the space spanned by three leading modes. (**c**) We designed the condition-fitted neural population model that lacks latent dynamics, and yet accurately reproduces oscillatory neural trajectories of the M1 dataset (Supplementary text 2). The plots show an example of the simulated firing rate (top panel) and the raster plot shows corresponding spiking activity (top panel) of this model for one reach. (**d**) Temporal covariance matrix of the model dataset for one reach direction (colorplot) is nearly Toeplitz. The first three latent variables (three-dimensional plot) are nearly sinusoidal curves of increasing frequencies. (**e, f**) Temporal covariance matrix of this model dataset for two opposite (e) or all eight reach directions (f) resembles the temporal covariance matrix of the M1 dataset (a, b). Neural activity projected in the space spanned by three leading modes traces oscillatory trajectories (three-dimensional plots) similar to those observed in real data (a, b).

To show this, we simulated a population of 84 neurons to match the number of neurons of the M1 center-out dataset. For this condition-fitted model, the simulated neurons were positioned in an eight-dimensional space with eight sources placed at unit vectors of this space. Each source is assigned to one reaching direction. The activity of the neurons for a reaching direction was determined by the Euclidean distance between the neuron’s location and this reaching direction source. n-th coordinate for each simulated neuron was obtained by taking the mean of a Gaussian function fitted to the activity of the corresponding recorded neuron for the n-th reaching direction (Supplementary text 2). This model does not rely on latent dynamics to generate the neural population activity since the neural activity results from the distance to the sources, irrespective of the location of other neurons. Therefore, the neural population activity is only dependent on the stimuli (Eq. 2). Alternatively, the same activity can be obtained with eight-dimensional wave propagating from one of eight different sources (Eq. 1). We used this model to simulate neural population activity datasets for one, two opposing, and all eight reach directions. In all three cases, temporal covariance matrix and the neural trajectory resembled those obtained from the M1 data (one direction: R^2^=0.84, CC=0.96; two opposing directions: R^2^=0.66, CC=0.90; all eight directions: R^2^=0.54, CC=0.87; Fig. 3c-f).

Consideration of the horseshoe effect described above may also help interpret recent results on long-term stability of recordings^8^, measures of the dynamical character of a system using tangling^9^, and the partition of neural activity into potent and null subspaces^32^ (Supplementary text 4-6).

Here, we showed that drawing conclusions about the latent dynamics of the system from the output of dimensionality reduction techniques applied to neural data can be misleading. We have established analytically that oscillatory trajectories are a necessary outcome of dimensionality reduction techniques applied on datasets with continuous variation in time, a rule otherwise known as the horseshoe effect. We then showed that models with and without latent dynamics can generate neural population activity with continuous variation in time that closely resembles recorded neural activity.

Numerous other studies described neural population activity with continuous variation in time in many different brain areas, across wide variety of tasks, and for different types of recordings, suggesting that this property is a ubiquitous feature of brain activity^2–7,9–11^. Reasons for which neural activity continuously varies in time is still debated, and could reflect a variety of processes, including variable time offsets between neurons and dynamics^33^, or propagating waves of neural activity^34^. Nonetheless, continuous variation in time is a feature of high-dimensional data in other science fields, where it is understood that horseshoe effect prevents drawing conclusions about dynamics of the system from the dimensionality reduction analysis results. Ultimately, it may be infeasible to identify the existence of the latent dynamics in high-dimensional data.

Considering the confound of the horseshoe effect, it is critical to complement dimensionality reduction analysis with adequate control analyses^8,35^ or analyses of other features in the data. For example, analysis of inter-conditional relationships may reveal latent organization of neural activity, independently of their dynamical nature (Supplementary text 3). Subsequently, uncovered features should always be verified by causal experimental manipulations.

## Acknowledgments

We thank Raeed H Chowdhury and Lee E. Miller for collecting the S1 dataset. We thank Matthew Smith for collecting the V4 dataset. TP is supported by the Swiss National Science Foundation (Grant #193542). The study was supported in part by the Swiss National Science

Foundation Ambizione grant to TM (PZ00P2_168103) and the Bertarelli Foundation Catalyst Fund (BC1709).

## Author contributions

T.P. and T.M. designed the study. M.G.P collected the M1 and PMd datasets. T.P. performed the analysis and modeling. T.P., M.G.P. and T.M. wrote the manuscript.

## Methods

All recording methods used in this article have been described in previous publications^27,28,30^. We here only briefly describe them.

## Experimental subjects

The animal neural datasets presented in this article were collected in accordance with the ethical standards of the Northwestern University and the University of Pittsburgh Institutional Animal Care and Use Committee. Data collection procedures were consistent with United States Federal guidelines. The neural data were recorded from three male Rhesus macaque (Macaca mulatta) monkeys.

## Behavioral tasks

For the M1, PMd, and S1 datasets, monkeys sat in a non-human primate chair and controlled a cursor displayed on a computer screen using a custom 2-D planar manipulandum. To successfully end a trial, the monkeys performed a standard center-out task – they started from the center and captured one of eight different 2 × 2 cm square targets that were evenly distributed on a circle of an 8 cm radius. We recorded the manipulandum handle position at a 1k-sample frequency. The behavioral task was controlled by a custom Simulink code (The Mathworks, Inc). The trial started when the monkeys moved the manipulandum to the center and captured the center target. Following a variable hold period (0.5 – 1.5 s), one of the eight outer targets appeared. After a variable instructed delay period (0.5 – 1.5 s), the monkeys received an auditory “go” cue, and the center target disappeared. The monkeys had to reach the outer target within one second, and had to hold the cursor there for another 0.5 s to successfully finish the trial.

For the V4 dataset, monkeys performed a fixation task while a sequence of static gratings and plaid stimuli were shown peripherally in a location centered on the V4 receptive fields. Each trial started with a period of fixation, followed by a sequence of ten flashed stimuli shown for 100 ms each and a new fixation period. The trial ended with a saccade to a random position target.

## Data acquisition

Monkeys were surgically implanted with microelectrode arrays (Blackrock Microsystems, Salt Lake City, UT), simultaneously in M1 and PMd for the first monkey, simultaneously in M1 and S1 for the second monkey, and in V4 for the third monkey. Blackrock Cerebus system (Blackrock Microsystems, Salt Lake City, UT) was used to record 96 channels of neural activity from each array. For the M1, PMd, and S1 datasets, the acquired data were processed using the Offline Sorter v3 software (Plexon, Inc, Dallas, TX) to identify action potential trains of individual neurons. There is a small possibility that duplicate neural activity can appear on different channels due to electrode shunting. We performed two procedures to ensure our data included only independent channels. First, we used Blackrock Cerebus system utility to identify and disable any channels with high crosstalk. Second, we computed the percent of coincident spikes between any two channels. This percentage was compared against an empirical probability distribution from all sessions of data. Any neurons with a coincidence above a 95% probability threshold were excluded from the analysis (approximately 15%–20% neurons per session). For the V4 dataset, data were sorted using customized semi-supervised clustering. Neurons that were not recorded stably throughout the session were excluded. For all datasets, we excluded neurons with trial-averaged firing rates below 1 Hz.

## Data processing

Neural spiking activity was counted in 10 ms bins. A smooth firing rate was then obtained for each electrode by convolving a Gaussian kernel (standard deviation 100 ms) with the binned spike train. For each trial of the center-out reaching task, we considered all time points between the go cue and the return to the initial start position after successful reaching of the target. Trials that were outside of the range of the median trial duration plus or minus 200 ms were discarded. To ensure that each trial contains the same number of samples (which is required for computing canonical correlation), we linearly interpolated reaching and returning sequence of each trial to match the number of samples found in the median trial duration. For the fixation task with visual stimuli, we used the full one-second length of the trials, shifted by 50 ms to account for the latency between the visual signal and the observed activity in V4. We used 124 trials of the same repeated stimulus sequence.

## Models

### Society for Neuroscience nationality model

We simulated the time on a set of 6 groups of neuroscientists from 6 distinct nationalities as Gaussian functions. Mean values of the Gaussians were uniformly spread between 0 and 1. Standard deviations and amplitudes were set to 0.02 and 0.1 for all Gaussians.

### Line neural population model

We simulated n neurons using a point-process generalized linear model with a log-link function and a conditionally Poisson distribution. The number of neurons was chosen to match the number of neurons of the data (100 for Fig. 1, 84 for the M1 dataset, 207 for the PMd dataset, 124 for the S1 dataset, and 26 for the V4 dataset). The dynamics of the instantaneous rate (also called conditional intensity function) of neuron *i*were modeled as:

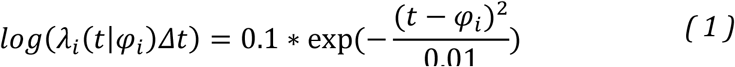

with the phase φ *i* ∈ 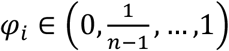 and the time t∈[0,1]. In this and all subsequent simulations, the number of time samples were chosen to match the number of samples of the data (396 for the M1 and PMd datasets, 74 for the S1 dataset, and 100 for the V4 dataset).

### Condition-fitted neural population model

We simulated 84 neurons. The dynamics of the instantaneous rate of neuron *i*were modeled by Eq. (1). The phase φ *ij* of each neuron *i*and target *j* is computed by fitting a Gaussian function to the firing rate of this neuron for this target. The center of the fitted Gaussian was then chosen as φ *ij* (Supplementary text 2).

### Measures

#### Canonical correlation

Canonical correlation finds linear transformations between two multidimensional trajectories such that the transformed trajectories are maximally correlated^8^. Formally, this corresponds to recursively defining a new basis (ξ *x*, ξ *Y*) for each multidimensional trajectory X and Y such that each of the dimension *i*∈ ⟦1, *N*⟧ of the basis maximizes the canonical correlation corr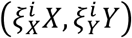 and is orthogonal to previously defined components.

#### Correlation between temporal covariance matrices

To assess correlation between two temporal covariance matrices, all values of one temporal covariance matrix were linearly regressed on all values of the other temporal covariance matrix. Correlation between two temporal covariance matrices was then quantified as the coefficient of determination R^2^ of this linear regression.

## Data availability

M1 and PMd data were used in previously published article^27^ and are available upon requests to the authors of this publication. S1 data are available on Dryad^28^. V4 data are available on CRCNS.org^31^.

## Code availability

Code needed to run the data processing, analysis and modeling is available at **DOI will be made available upon acceptance**.

