## Supplemental file for "Misinterpreting the horseshoe effect in neuroscience"

#### Supplementary material

Timothée Proix, Matthew G. Perich, Tomislav Milekovic

### Extended Data Figures

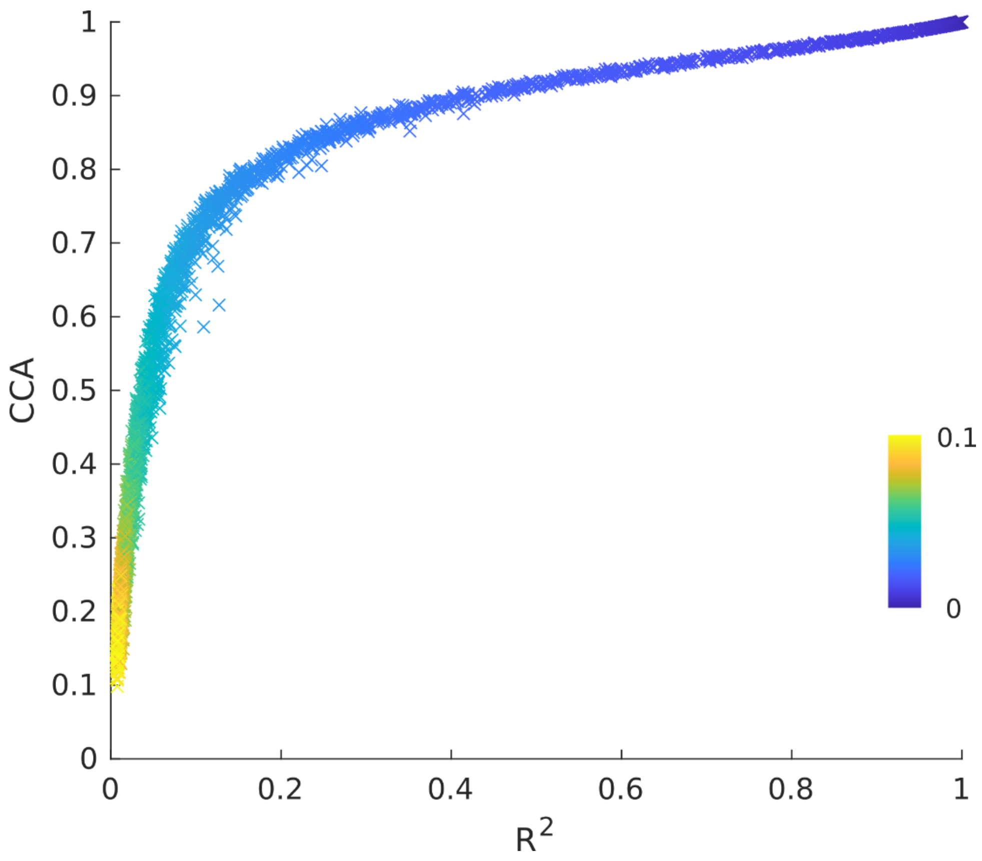


***Extended Data Fig. 1. The Pearson's correlation measure $R^2$, calculated between the neural dataset temporal covariance matrix and the ideal Toeplitz matrix, is proportional to the canonical correlation (CC), calculated between the four-dimensional neural trajectory and the four-dimensional horseshoe curve.*** *To assert that deviations from the horseshoe curves are proportional to the similarity of the temporal covariance matrix to the Toeplitz matrix, we computed the Pearson's correlation $R^2$ and the canonical correlation CC for different levels of white noise ($\mathcal{N}(0,\sigma)$ with $\sigma$ between 0 and 0.1 indicated by*

*the colormap) added to the time series simulated with the line model (Eq. (1), Fig. 1e-f). Each dot on the scatterplot corresponds to a simulation of the line model with a given level of white noise added to the resulting time series. We performed 10 simulations for each noise level. The unveiled relationship between* $R^2$ *and CC is approximatively proportional, monotonic and bijective.*

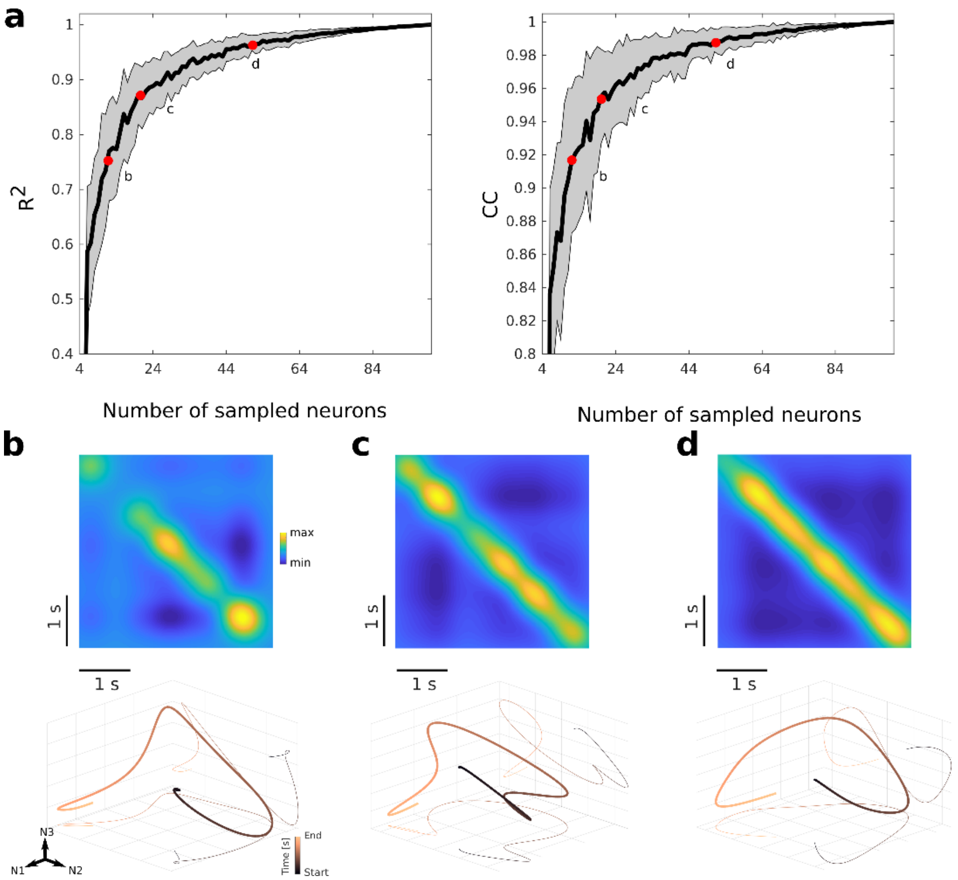


***Extended Data Fig. 2. Random partial sampling of the neural population that approximately exhibits continuous variation in time has only a minor impact on the shape of the latent oscillatory trajectory.*** *To demonstrate this feature, we selected random subpopulations of the line model population comprising 100 neurons (Fig. 1d-e). For each selection, we computed temporal covariance matrix and three-dimensional neural trajectories using PCA. We then computed* $R^2$ *and CC against the Toeplitz temporal covariance matrix and the horseshoe curve, respectively. We performed 100 random selections for each subpopulation size, from 1 to 100. We found that such random partial sampling only had a minor impact on the shape of the oscillatory trajectory if the sampling is performed with more than a few neurons. (**a**) Left plot shows Pearson's correlation* $R^2$ *between the temporal covariance matrix of the entire neural population, which has the Toeplitz structure (Fig. 1f), and the randomly selected subpopulations. Right plot shows canonical correlation CC between the three-dimensional neural trajectories of the entire neural population, which is a horseshoe curve (Fig. 1f), and randomly selected subpopulations. (**b**) Temporal covariance matrix (up) and three-dimensional neural trajectory (down) of a representative dataset for random partial sampling with ten neurons. The representative dataset was selected by taking a dataset*

*with a median $R^2$. $R^2$ and CC values of this dataset are shown in (a) by a red dot. (**c**) Same as (b) for 20 neurons. (**d**) Same as (b) for 50 neurons.*

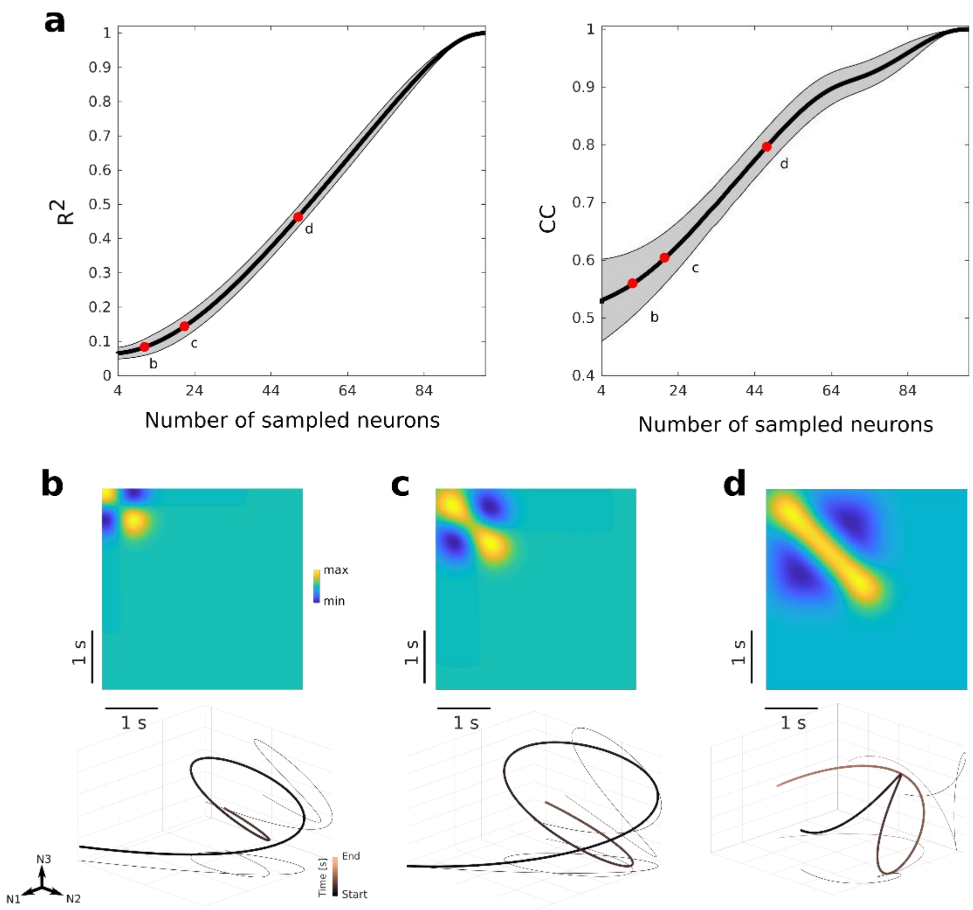


***Extended Data Fig. 3. Biased sampling of the neural population that exhibits continuous variation in time has a limited impact on the shape of the three-dimensional neural trajectory.*** *To demonstrate this feature, we selected biased subpopulations of the line model population comprising 100 neurons. We performed biased selections by only selecting a number of successive neurons on the line, where this number is the subpopulation size. We systematically renewed the operation, progressively shifting the selected neurons. Experimentally, bias sampling can occur if neurons exhibiting continuous variation in time are locally tuned, and neural recordings are only performed on a spatially localized subset. For each selection, we computed temporal covariance matrix and three-dimensional neural trajectories using PCA. We then computed* $R^2$ *and CC against the Toeplitz temporal covariance matrix and the horseshoe curve, respectively. We performed 100 biased selections for each subpopulation size, from 1 to 100. We found that such random partial sampling had a limited impact on the shape of the oscillatory trajectory, with the deviations more pronounced than for random partial sampling (Extended Data Fig. 2). (**a**) Left plot shows Pearson's correlation* $R^2$ *between the temporal covariance matrix of the entire neural population, which has the Toeplitz structure (Fig. 1f), and the biased subpopulations. Right plot shows canonical correlation*

*CC between the three-dimensional neural trajectories of the entire neural population, which is a horseshoe curve (Fig. 1e), and biased subpopulations. (**b**) Temporal covariance matrix (up) and three-dimensional neural trajectory (down) of a dataset of ten selected neurons. The representative dataset was selected by taking a dataset with a median* $R^2$*.* $R^2$ *and CC values of this dataset are shown in (a) by a red dot. (**c**) Same as (b) for 50 selected neurons. (**d**) Same as (b) for 100 selected neurons*

### Supplementary text

#### 1. Analytical derivation of the horseshoe effect

##### Latent variables obtained after applying principal component analysis to a neural dataset are proportional to the eigenvectors of the temporal covariance matrix

Let's assume that our centered data matrix X is a $n$ x $p$ matrix, with $n$ the number of observations (or time), and $p$ the number of neurons. Principal component analysis (PCA) applied to this dataset finds the modes (also called principal component coefficients) and latent variables (also called scores). Those can be obtained by computing the eigenvectors $v^k$ and eigenvalues $\lambda_k$ of the covariance matrix $X^T X$ of the dataset (we removed normalization factor for simplicity, without loss of generality) with $k \in [\![1, p]\!]$, i.e.

$$X^T X v^k = \lambda_k v^k \quad (2)$$

The PCA modes are then the eigenvectors $v^k$ of the covariance matrix $X^T X$, and the latent variables $Xv^k$ are the product of the centered data $X$ with the eigenvectors $v^k$.

To obtain an expression for the latent variables, we use the so-called 'PCA transpose trick': we multiplied both side of *Eq. 2* by $X$:

$$X^T X v^k = \lambda_k v^k$$

$$XX^T X v^k = \lambda_k X v^k$$

$$XX^T w^k = \lambda_k w^k \quad (3)$$

with $w^k = Xv^k$ the $k$-th eigenvector of the transposed covariance matrix $C = XX^T$. Thus, the eigenvalues are the same for the covariance and the transposed covariance matrix. Further, the eigenvectors of the transposed covariance matrix are the latent variables of the PCA applied on the centered data $X$. For data evolving in time, with $n$ observations of the neuronal firing rate of $p$ neurons, the transposed covariance matrix $C$ is called the temporal covariance matrix. Therefore, the eigenvalues of the temporal covariance matrix can be derived using the previous formula:

$$Cw^k = XX^T w^k = \lambda_k w^k \quad (4)$$

Note that the PCA latent variables (*Eq. 3*) are therefore identical to the eigenvalues of the temporal covariance matrix (*Eq. 4*).

#### Eigenvectors (PCA latent variables) of the temporal covariance matrix form a Fourier series

Let the temporal covariance matrix be circulant. This matrix is obtained when the dataset has continuous variations in time with periodic boundary conditions (i.e. when the continuous variation in time is maintained between the last and first data points). Such dataset is obtained from our SfN example (Fig. 1) if the corridor is circular. The eigenvalues and eigenvectors of a circulant matrix can be shown to have a very specific form, as follows[1]. We note $C = XX^T$ the temporal covariance matrix. The eigenvalue $\lambda_k$ and eigenvector $w^k$ problem is a system of $n$ equations:

$$Cw^k = \lambda_k w^k \qquad (5)$$

C is a circulant matrix

$$\begin{bmatrix} c_0 & c_1 & c_2 & & \cdots & c_{n-1} \\ c_{n-1} & c_0 & c_1 & c_2 & & \vdots \\ & c_{n-1} & c_0 & c_1 & \ddots & \\ & \ddots & \ddots & \ddots & & c_2 \\ \vdots & & & & & c_1 \\ c_1 & \cdots & & c_{n-1} & c_0 & c_0 \end{bmatrix}$$

and thus each entry can be described using only a single index:

$$c_{l,m} = c_{(m-l)\,\mathrm{mod}\,\mathrm{n}}$$

System (3) can thus be written as a system of n difference equations, for $j = 0, 1, \ldots, n-1$:

$$\sum_{l=0}^{j-1} c_{n-j+l}\, w_l^k + \sum_{m=j}^{n-1} c_{m-j}\, w_m^k = \lambda_k w_j^k$$

Changing the summation dummy variables to o= $n - j + l$ and $q = m - j$ gives

$$\sum_{q=0}^{n-1-j} c_q\, w_{q+j}^k + \sum_{o=n-j}^{n-1} c_o\, w_{o-(n-j)}^k = \lambda_k w_j^k$$

To find a solution to this system of equations, we make an ansatz and then prove that it solves this system. Our ansatz for the eigenvector is $w_j^k = \frac{1}{\sqrt{n}} e^{-2\pi ijk/n}$, i.e.

$$w^k = \frac{1}{\sqrt{n}}(1, e^{-2\pi ik/n}, \ldots, e^{-2\pi ik(n-1)/n})'$$

which yield:

$$\sum_{q=0}^{n-1-j} c_q e^{-2\pi iqk/n} e^{-2\pi ijk/n} + \sum_{o=n-j}^{n-1} c_o\, e^{-2\pi iok/n} e^{2\pi ik} e^{-2\pi ijk/n} = \lambda_k e^{-2\pi ijk/n}$$

$$\sum_{q=0}^{n-1-j} c_q e^{-2\pi iqk/n} + \sum_{o=n-j}^{n-1} c_o\, e^{-2\pi iok/n} = \lambda_k$$

$$\lambda_k = \sum_{l=0}^{n-1} c_l e^{-2\pi ilk/n}$$

These eigenvalues/eigenvectors pairs solve the equation (3), for all $k$, completing the proof.

We now look for real eigenvectors of equation (3). If $\lambda_k$ and $w^k$ are eigenvalues and eigenvectors of (3), then $\lambda_{-k}$ and $w^{-k}$ are eigenvalues and eigenvectors as well, and

$$\lambda_{-k} = \sum_{l=0}^{n-1} c_l e^{-2\pi il(-k)/n}$$

Changing the dummy variable m=n-l

$$\lambda_{-k} = \sum_{m=1}^{n} c_{n-m} e^{-2\pi i(n-m)(-k)/n}$$

If the temporal covariance matrix is symmetric, then $c_m = c_{n-m}$, and then

$$\lambda_{-k} = \sum_{m=1}^{n} c_m e^{2\pi ik}\, e^{-2\pi imk/n}$$

$$\lambda_{-k} = \sum_{m=1}^{n-1} c_m\, e^{-2\pi imk/n} + c_0$$

$$\lambda_{-k} = \sum_{m=0}^{n-1} c_l e^{-2\pi imk/n} = \lambda_k$$

Using this property, we then look for real eigenvectors:

$$\frac{1}{2}\left(\lambda_k w^k + \lambda_{-k} w^{-k}\right) = \lambda_k \left(\frac{w^k + w^{-k}}{2}\right) = \frac{1}{2}\left(Cw^k + Cw^{-k}\right) = C\left(\frac{w^k + w^{-k}}{2}\right)$$

The eigenvectors $\frac{w^k+w^{-k}}{2} = \frac{1}{\sqrt{n}}(1, \cos(\frac{2\pi jk}{n}), \ldots, \cos(\frac{2\pi(n-1)k}{n}))$ are real. Similarly, $\frac{w^k-w^{-k}}{2} = \frac{1}{\sqrt{n}}(1, \sin(\frac{2\pi jk}{n}), \ldots, \sin(\frac{2\pi(n-1)k}{n}))$ are real eigenvector for the eigenvalue $\lambda_k$. Thus the eigenvectors of the temporal covariance matrix are the Fourier series, as well as the latent variables $Xv^k = w^k$ obtained from PCA.

##### Eigenvectors of Toeplitz temporal covariance matrix are approximate Fourier series

In the more realistic case where the temporal covariance matrix is Toeplitz and not circulant, analytical results can be obtained if the Toeplitz matrix is symmetric, which is always the case for covariance matrices. In particular, it can be shown that the asymptotic eigenvectors of the symmetric Toeplitz matrix converge to Fourier modes with some edge corrections[2].

#### 2. Condition-fitted neural population model

The condition-fitted neural population model is obtained by fitting, for each neuron $i$ and each reaching direction $j$, a Gaussian to the firing rate $r_{ij}$ of the corresponding neuron for that reaching direction:

$$\mathcal{N}(\mu_{ij}, \sigma, A_{ij}) = \frac{A_{ij}}{\sigma\sqrt{2\pi}}\, e^{-(r_{ij}-\mu_{ij})^2/2\sigma^2}$$

where $\mu_{ij}$ was the mean, σ standard deviation, and $A_{ij}$ amplitude of the Gaussian. Mean $\mu_{ij}$ and amplitude $A_{ij}$ were fitted by minimizing the Euclidean distance between the firing rate of the neuron $i$ for each reaching direction $j$ and a Gaussian function. Because we here showcase a simple model with approximate continuous variation in time, we only used the fitted Gaussian means as parameters to simulate the activity, while the amplitude and standard deviation of all Gaussians were set to $0.1 \cdot \sqrt{0.005 \cdot 2\pi}$ and $\sqrt{0.005}$, respectively. Thus, the activity of neurons is given by:

$$log(\lambda_{ij}(t|\mu_{ij})\Delta t) = 0.1 * \exp(-\frac{(t-\mu_{ij})^2}{0.01})$$

A useful representation of this model can be constructed in an eight-dimensional space. We can assign an “activity source” for each of the eight reaching directions as a location in this 8D space (Supplementary Fig. 1). We can place these sources at the space unit vectors, i.e. $s_1 = (1, 0, 0, 0, 0, 0, 0, 0)$ for the source assigned to the first reaching direction, $s_2 = (0, 1, 0, 0, 0, 0, 0, 0)$ for the second reaching direction source, etc. We can then place a neuron in that space so that the Euclidean distance between this neuron’s location $x_i$ and each source $s_j$ correspond to the fitted Gaussian means for this neuron:

$$d(x_i, s_j) = \|x_i - s_j\|_2 = \mu_{ij}$$

It can be ensured that at least one solution exists by linearly warping Gaussian centers to the interval $[\sqrt{2}, \frac{3}{2}\sqrt{2}]$. In this eight-dimensional functional space, neurons' firing rate can be intuitively understood either as a delayed activity of the source activity, with the delays equal to the Euclidean distance in this functional space, or as the activity resulting from a wave propagating from that source. Note that the former does not constitute a dynamical system, while the latter does. Each source sets the timing of the activity of other neurons once the reach in that direction has been initiated. We call this space "functional" because Euclidean distances between neurons and sources do not represent actual physical distances, but rather the similarity between the activity patterns of neurons across different reaching directions.

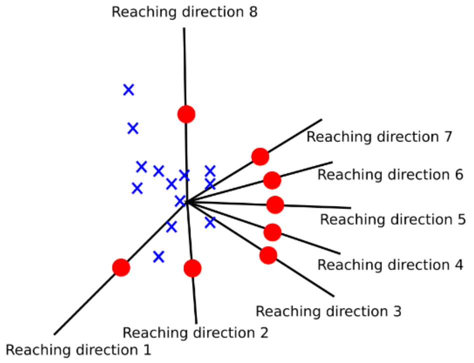


***Supplementary Fig. 1. Condition-fitted neural population model.*** *This model comprised a population of 84 neurons fitted to the neural recordings. For one of eight reaching directions, activity of each neuron resulted from the distance to the source assigned to that reaching directions. The eight sources were placed on the unit vectors of the eight-dimensional space (red points).*

#### 3. Reaching directions are spatially organized in a reduced functional space

We showed that the condition-fitted model, which arranges neurons and sources in a multidimensional functional space, can accurately reproduce the activity of a neural population across several conditions (Fig. 3f). To better understand the relationship between sources, we looked for lower dimensional representations of the functional space. In a functional space with the dimensionality equal to the number of sources, the sources can be entirely uncorrelated – as was the case with our condition-fitted neural population model. Yet, in a functional space with the dimensionality lower than the number of sources, the sources become correlated. If the sources are placed to accurately reproduce recorded neural population activity, the correlations between sources will highlight their functional similarities. To show this, we designed an optimization algorithm to place neurons $\tilde{x}_i$ and sources $\tilde{s}_j$ in an N-dimensional space, where N is less or equal to the number of conditions. In this space, the distance between each neuron and each source is optimized to be proportional to the mean of the Gaussian fitted to that neuron's activity recorded during that condition.

To generate this "reduced" model, we designed a cost function that computed the L2 norm between the Gaussian means $\mu_{ij}$ and the Euclidean distances between each neuron and each source:

$$\mathcal{C}(\tilde{x}_i, \tilde{s}_j) = \left\| \mu_{ij} - \left\| \tilde{x}_i - \tilde{s}_j \right\|_2 \right\|_2$$

We here briefly describe the algorithm steps:

1. Choose the dimensionality of the reduced space N
2. For n_ini different initial positions
    a. Initialize source and neuron positions by randomly drawing from a standard Gaussian distribution.
    b. For m_ite iterations
        i. For each source *j*
            1. For each dimension of the reduced space, move this source along each of the current dimension by a large positive and large negative spatial step.
            2. Calculate the cost function for each of the newly obtained positions.
            3. If the cost function of at least one of the new positions is smaller than the cost function of the previous position, keep the source location with the smallest cost function.
            4. Else repeat steps 1-4 with a halved spatial step, until a new minimum is found (under a given spatial threshold, this is simply chosen as the old position).
        ii. For each neuron $i$
            1. For each dimension of the reduced space, move this neuron along each of the current dimension by a large positive and large negative spatial step.
            2. Calculate the cost function for each of the newly obtained positions.

3. If the cost function of at least one of the new positions is smaller than the cost function of the previous position, keep the neuron location with the smallest cost function.
4. Else repeat steps 1-4 with a halved spatial step, until a new minimum is found (under a given spatial threshold, this is simply chosen as the old position).

We chose n_ini = 100 to ensure that a large sample of the cost space is covered, and m_ite=100 to ensure that the algorithm converged.

The full code of the algorithm is provided at **#URL WILL BE PROVIDED UPON ACCEPTANCE.**

We verified that the algorithm converged by computing the cost as a function of the number of iterations across all initial conditions. We found that the algorithm was converging in tens of iterations for all dimensions considered, and that similar values of the cost functions were reached across different initializations (Supplementary Fig. 2a). At each iteration of the algorithm, we also computed CC values between the positions of the sources across algorithm runs for different initial conditions. For instance, at iteration n, we computed CC values between the source positions for initializations 1 and 2, between initializations 1 and 3, 2 and 3, etc. This gave us a distribution of CC values for each iteration of the algorithm (Supplementary Fig. 2b). We found that CC values tended towards a value of 1 already after tens of iterations. This result shows that the source positions for different initializations of the optimization algorithm are rapidly converging.

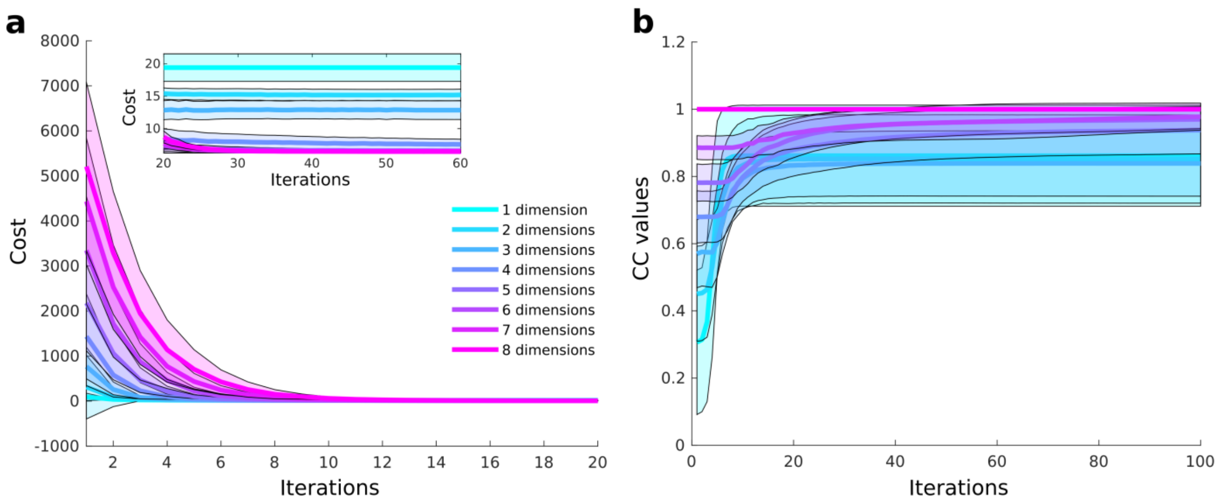


***Supplementary Fig. 2. Solutions found by the optimization algorithm are consistent across random initializations.*** *(**a**) Main panel plot shows the median cost as a function of the number of iterations for up to 20 iterations for different dimensionality of the model. Inset panel plot shows the same cost for iterations 20 to 60. Colored transparent tubes show the standard deviation. (**b**) The plot shows median CC values between positions of the sources across different iterations as a function of the number of iterations. Colored transparent tubes show the standard deviation.*

We systematically computed $R^2$ and CC values as a function of the dimensionality across all initializations of the algorithm (Supplementary Fig. 3). Both $R^2$ and CC median values across initializations increase as a function of dimensionality. Note that, as opposed to the eight-dimensional space constructed arbitrarily for the condition-fitted neural population model with a perfect representation of the fitting, here we build an approximation automatically, which explain why we don't reach the $R^2$ and CC values reported in the main text for eight dimensions. Thus, a large part of the neuronal activity can be accounted for by a low dimensional functional space, whose dimensionality can be chosen for instance from the $R^2$ and CC plot.

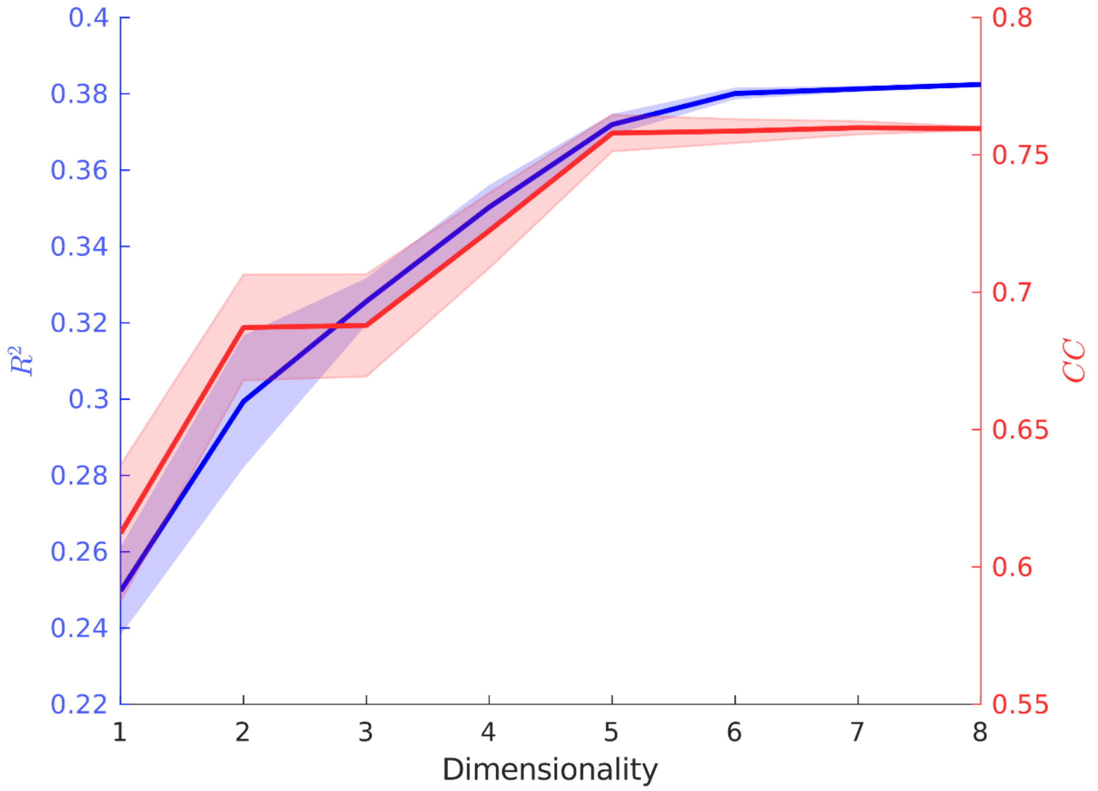


***Supplementary Fig. 3. Model fit to neuronal recordings improves as a function of the dimensionality of the reduced functional space.*** *For each number of dimensions, we computed $R^2$ values (blue, left y-axis) between the temporal covariance matrix generated by the reduced model, and the temporal covariance matrix of the full dataset. We computed the CC values (red, right y-axis) between neural trajectories generated by the reduced model and neural trajectories from the full dataset. Full lines and shaded areas show the median and standard deviations across all initializations of the optimization algorithm.*

We examined the three-dimensional representation for the initialization of the algorithm with the highest $R^2$ (Supplementary Fig. 4). Although initiated randomly, the sources' optimal positions were organized in a circular arrangement: the sources were approximately placed on a tilted plane and sources for neighboring reaches were closer compared to dissimilar reaches. Thus, the algorithm recovered two main features of the neural recordings (Fig. 3b), which can only be accounted for by at least three dimensions:

1. Circular symmetry of the sources: neural activity of reaching targets with small angular difference are more similar to each other than between reaching target with large angular difference. This can be observed in the temporal covariance matrix, with high correlation values distributed along

the between-condition block diagonals for pairs of similar reaching directions. This can also be observed in the similarity of three-dimensional neural trajectories for similar reaching directions.

2. Quasi-orthogonality of the sources for opposite reaching direction: neuronal activity for two opposite reaching directions are very dissimilar. This is reflected in the temporal covariance matrix, where between-condition blocks for opposite reaching directions do not have large sequence of correlation. Furthermore, three-dimensional neural trajectories of opposite reaching directions are quasi-orthogonal.

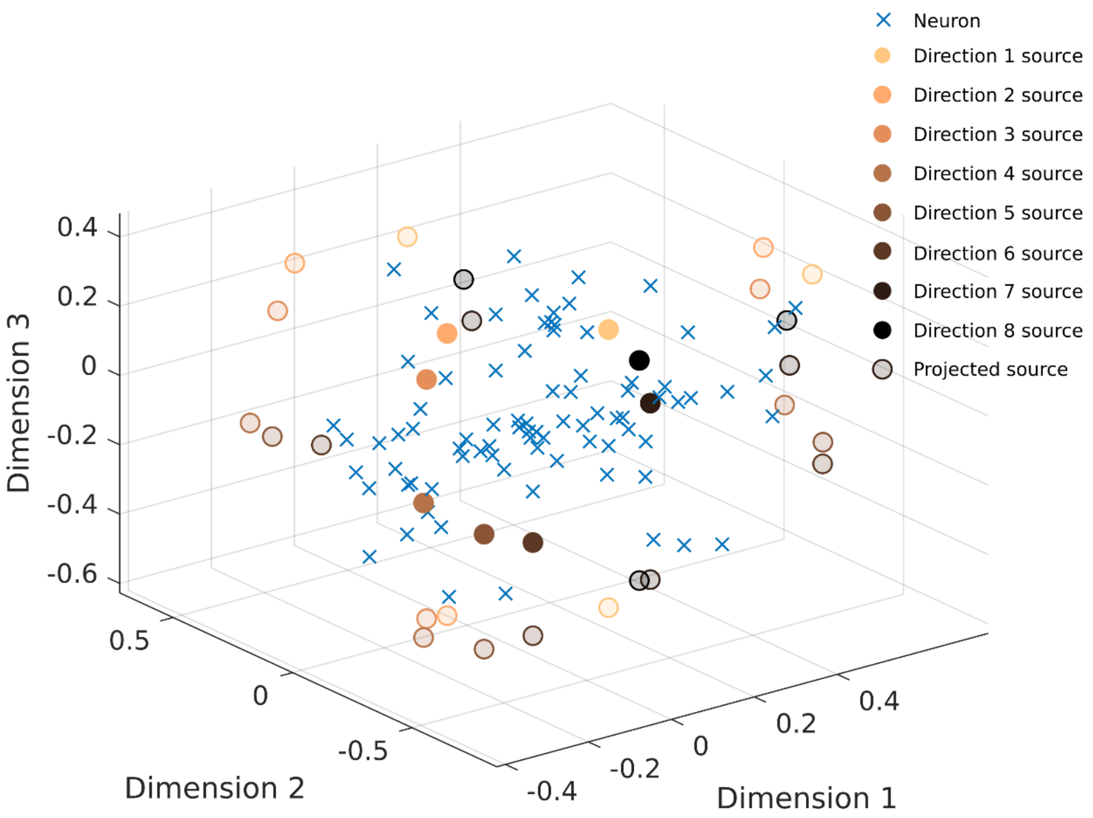


***Supplementary Fig. 4. Reduced functional space in three dimensions.*** *Source locations for each reaching direction are shown as opague spheres. Transparent spheres show the source locations projected on planes spanned by two of the three dimensions. The color of the spheres represents the reaching direction. Sources are organized in a circular arrangement, and ordered by reaching direction. Neuron locations in the functional space are shown as blue crosses.*

#### 4. Long-term stability of neural trajectories is expected due to the horseshoe effect

A recent study described stability of neural population activity recordings collected over up to two years[3]. In this study, authors trained six monkeys to perform a center-out reaching task while recording neural activity with a Utah array implanted in M1, S1 or PMd brain regions. The task was repeated many times across different days, spanning time duration up to two years. When applying PCA on neural recordings from different days made by the same Utah array[3], the contribution of neurons in the leading modes changed over time. However, the three-dimensional neural trajectory preserved its shape that was easily matched between days by a linear transformation. The shape of the trajectory was similar to the horseshoe curve. The measure used to assess the correlation between the neural trajectories was the canonical correlation.

As elaborated in the main text, due to the horseshoe effect, the leading latent variables, which explain the largest portion of the variance of the neural population activity, have oscillatory trajectories. These oscillatory trajectories are expected as long as the firing of the neurons remains sequential, and are little affected by the changes in the firing rate statistics or by the evolution of the waveform shapes of the neural action potentials that are typical for the longitudinal Utah array recordings. Therefore, high CC values across time obtained in that study are mainly accounted for by the similarity of the oscillatory trajectories, which remain trivially similar due to the horseshoe effect.

To a lesser degree, CC values are also affected by the spatial ordering of the trajectories corresponding to different targets, in the case of this study the reaching directions. Stable high CC values thus also show that the similarity of the neural activity between conditions remains stable across days. Unlike the oscillatory shape of the neural trajectories for individual targets that are generated by the horseshoe effect, this feature of the data, here hypothesized to arise from a functional space (Supplementary text 3), is not trivial and reflects important structure of the neural activity preserved over time.

The importance of the functional space for low dimensional trajectories is further illustrated by another recent study analyzing M1 neural population activity recorded by Utah arrays implanted in monkeys as they performed different groups of upper-limb tasks[5]. These task groups included standard isometric center-out force (applying force in center-out directions without movement) and center-out movements, as well as reaching and grasping balls of different sizes. Oscillatory trajectories were computed by performing PCA on all concatenated trials and targets of each task. CC-values between trajectories for different tasks were lower than the CC-values between trajectories for the same task recorded on different days. This result is in line with our functional space hypothesis. Ordering of neural trajectories corresponding to different targets is specific to each task, thus reflecting the task-specific structure of the neural activity. Even though tasks may have similar structure (e.g. center-out movement group versus isometric center-out force), the differences between the trajectories will be larger than when comparing the trajectories of the same task derived on different days.

Interestingly, the functional structure of a task may be encoded in a similar way across different animals of the same species reflecting, for example, the invariant way that the neural populations encode the same behavioral strategy shared among the animals. The invariance of that functional space would result in a similar shape of neural trajectories for that task across the animals, as long as the neural trajectories are derived from neural population activity of the same brain regions. CCA between these two sets of neural trajectories will provide a mapping from the low dimensional space of one animal to the other. Interestingly, neural decoders calibrated on neural population activity of one animal can also be mapped to another animal while maintaining most of its accuracy. Results from two recent studies already support this hypothesis. The first study shows that neural decoders indeed generalize across neural population activity recorded in different monkeys[4]. The second utilized this property to calibrate a neural decoder from M1 neural population activity recorded in one monkey to detect reach and grasp attempts of another monkey. These detected attempts were used to trigger spinal cord stimulation in the other monkey that restored its ability to reach and grasp after the paralyzing spinal cord injury[6]. This demonstration illustrated that invariance of the functional space may be utilized for transfer of neural decoders in neuroprosthetic applications. Nonetheless, further research is needed to characterize the similarity and differences of functional spaces between the animals.

#### 5. Structure of the functional space interacts with the horseshoe effect to affect tangling, independently of the dynamical character of a system

The horseshoe effect can affect measures based on dimensionality reduction results and, therefore, confound the interpretation of these measures. Specifically, recent studies proposed to quantify the dynamical character of a system using a measure called tangling[7,8]. Tangling evaluates if close-by low-dimensional neural trajectories, obtained by applying PCA to neural population recordings, follow similar gradients:

$$Q(t) = \max_{t'} \frac{\|\dot{x}_t - \dot{x}_{t'}\|^2}{\|\dot{x}_t - \dot{x}_{t'}\|^2 + \varepsilon}$$

with $x_t$ the neural state at time $t$, $\dot{x}_t$ its temporal derivative, and $\varepsilon$ a small constant preventing zero division. Low tangling values are then interpreted as evidence for the existence of a latent dynamical system. For example, applying PCA to EMG recordings from a task involving forward and backward cycling resulted with trajectories rotating in opposite directions and, thus, in high tangling values. Applying PCA to the M1 neural population recordings from the same task resulted with co-rotational trajectories and, thus, with much lower tangling values[7]. Authors interpreted these low tangling values as evidence for latent dynamics driving the M1 neural population activity, while high tangling values of the EMG recordings were interpreted as signature of a predominantly input-driven system.

To understand if the horseshoe effect influences tangling results, we repeated the same analysis with two models, each with a different functional space structure. The first model was our line model with 100 simulated neurons (Fig. 1e-f; Supplementary Fig. 5a-b). The dynamics of the instantaneous rate of neuron $i$ were modeled by *Eq. 1*. We modeled two reaching directions $j$, represented as sources placed at each extremity of the line of neurons. The phase $\varphi_{ij}$ is calculated as the distance between the position $x_i$ of each neuron in a one-dimensional space and the position $s_j$ of the source, which is different for each reaching direction:

$$\varphi_{ij} = \left\|x_i - s_j\right\|_2$$

For neurons, $x_i \in [\![1{,}100]\!]$. For sources, $s_1 = 1$, $s_2 = 100$.

For the second "square" model, we simulated 100 neurons, arranged on a square (Supplementary Fig. 5c-d). The dynamics of the instantaneous rate of neuron $i$ were modeled by *Eq. 1*, and the phase as:

$$\varphi_i = \|x^T - s^T\|_2 / \max_i (\|x^T - s^T\|_2)$$

Each neuron *i* and source *j* were given as a set of two Cartesian coordinates $x = (x_i, y_i)$ and $\left(s_j^x,\, s_j^y\right)$, with the superscript T denoting the transpose operator. For neurons, $x_i \in [\![1{,}10]\!]$, and $y_i \in [\![1{,}10]\!]$. For sources, $s_1 = (5.5, 1)$, and s $s_2 = (5.5, 10)$.

Neither of these models constitute a dynamical system, although similar activity can be generated with a dynamical system (e.g. by simulating a wave propagating from each source). For each model, we then concatenated the two simulated reaching directions and performed PCA. For the line model, neural trajectories for two reach directions were exactly matching, but rotating in opposite directions. This is because for the line model, the number of neurons activated at each time point was exactly the same across time and for both reaching directions, resulting in block temporal covariance matrix where each block has a Toeplitz structure, and exactly opposite oscillatory trajectories generated by the Horseshoe effect after dimensionality reduction. This two overlapping counter-rotating oscillatory trajectories result in high tangling values, because of the large difference between temporal derivatives of each trajectory (Supplementary Fig. 5e). In contrast, the square model neural trajectories are not overlapping, and were rotating in the same direction, resulting in low tangling values (Supplementary Fig. 5f). This is because the number of neurons activated at each time point is no longer the same across time, resulting in deviations of the blocks of the temporal covariance matrix from the Toeplitz structure. These deviations are reflected in departure of the neural trajectories for each reaching direction from the Fourier series. The underlying functional space of the system, independently of the dynamical character of the system, interacts with the horseshoe effect to generate the neural trajectories and, therefore, affect the resulting tangling values.

Coming back to the previous comparison of tangling of EMG and neural population recordings, the difference in tangling between the two systems may have been affected by the difference in the functional space underlying the two recorded systems. EMG recordings were collected from eight hand and arm muscles. Their sequential activation may constitutes a low dimensional functional space. In contrast, neural activity dataset was recorded from the motor cortex, which may constitute a higher dimensional functional space. Even though the analysis was made on a low-dimensional trajectory that was matched in dimensionality between the two datasets, as we imitated here with our line and square models, the underlying functional space of the two datasets through an interaction with the horseshoe effect may have affected the tangling values and, therefore, may have influenced the interpretation of their difference.

Nonetheless, our modeling example only highlights that the functional space interact with the horseshoe effect to affect the tangling. Further research is needed to determine the underlying functional space of a dataset, and to quantify the effect of different functional spaces on tangling. One possible way to identify the functional space is to utilize the functional space optimization algorithm described in this study (Supplementary text 3).

Note that the effect described here will not affect the comparison of tangling values calculated on datasets recorded from the same system. For example, comparing tangling of the M1 recordings performed on different days or during different phases of the experiment will remain unaffected.

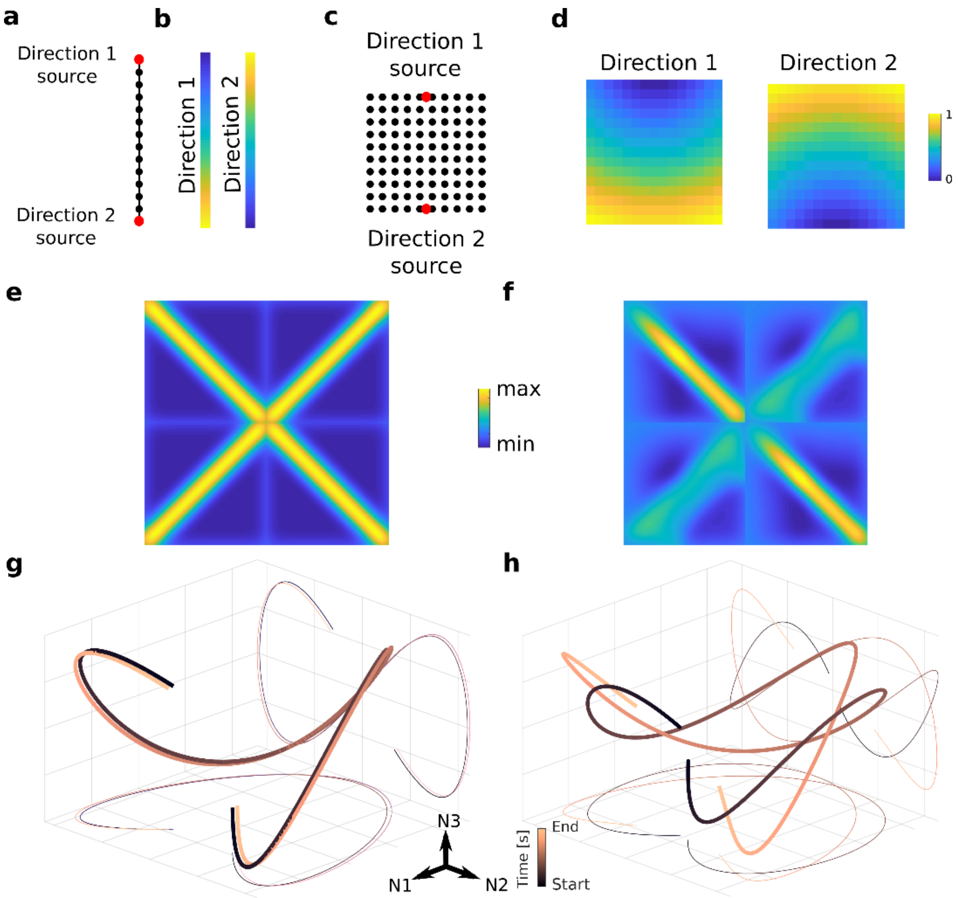


***Supplementary Fig. 5. Oscillatory trajectories change when changing the functional structure of the system, regardless of the dynamical nature of the system.*** *(**a**) Schematic of the line model. Red points indicate the position of the sources. (**b**) Values of the phase shift for simulated neurons of the line model. (**c**) Schematic of the square model. (**d**) Values of the phase shift for simulated neurons of the square model. Neither the line nor the square model incorporates latent dynamics. (**e**) Temporal covariance matrix for the line model and two reaching directions. (**f**) Temporal covariance matrix for the square model and two reaching directions. (**g**) Plot shows the three-dimensional neural trajectory of the line model dataset for two reach directions concatenated. The curves for two reach directions overlap each other (a slight spatial shift was added for visualization), but move in the opposite directions. (**h**) Plot shows the three-dimensional neural trajectory of the square model dataset for two reach directions concatenated. The curves for two reach direction cross but do not overlap, and move in the same direction.*

#### 6. Horseshoe effect can result in fortuitous generation of potent and null subspaces

Recent studies argued that neural modes of the activity of two regions can be partitioned into subspaces where information is shared across the two regions (called potent space) and subspaces for which activity of one region has no effect on the other region (called null space)[9,10]. These subspaces have been thought to offer a mechanism by which information is shared across brain regions and from a brain region to a downstream target, including muscles. However, the horseshoe effect can lead to trivial identification of the potent and null spaces regardless of the relationships between the two systems (e.g. cortical regions or muscles) that the datasets were recorded from. To illustrate this, we used the M1 and PMd center-out dataset (Fig. 2a-h). Dimensionality reduction of M1 and PMd neural recordings yielded four leading latent variables that approximately followed the Fourier series (Supplementary Fig. 6a). We then computed the potent and null spaces of the first four PMd neural modes to explain the first two neural modes of the M1 space (Supplementary Fig. 6b). The potent space largely aligned with the first two PMd neural modes and the null space with the modes three and four.

To understand why the horseshoe effect can trivially generate this relationship, we simulated neural activity in M1 and PMd using our 8D condition-fitted neural population model for one reaching direction. Simulated neural activity of each brain regions exhibits approximate continuous variation in time and both have Toeplitz-like temporal covariance matrices, even though there is no connection between the two regions. Due to the horseshoe effect, the latent variables of each region then trace the Fourier series at a particular frequency (Supplementary Fig. 6c). The subspace formed of the first two PMd latent variables, $[\cos(2\pi\, t/T), \sin(2\pi\, t/T)]$, completely explain the subspace formed of the first two M1 latent variables oscillating at the same frequency. The following PMd latent variables, $[\cos(4\pi\, t/T), \sin(4\pi\, t/Tt), \cos(6\pi\, t/T), \dots]$, are necessarily orthogonal to the first two M1 latent variable, $[\cos(2\pi\, t/T), \sin(2\pi\, t/T)]$. Therefore, the first two PMd modes form the potent space for the first two M1 modes, while the rest of the PMd dataset forms the null space, irrespective of any relationship between the cortical regions (Supplementary Fig. 6d). This example shows that null and potent spaces can be derived even between two disconnected brain regions as long as the neural population activity of both regions approximately exhibit continuous variation in time. In other words, while the neural activity of the brain regions themselves do not necessarily correlate with each other, the resulting principal components of those neuronal activities do correlate because of the horseshoe effect. Yet, as stated by the well-known aphorism, correlation is not causation. Therefore, the mere existence of potent and null subspaces cannot be taken as evidence of information exchange across brain regions, since these relationships can emerge from a common feature of neural cortical activity reflected in both regions.

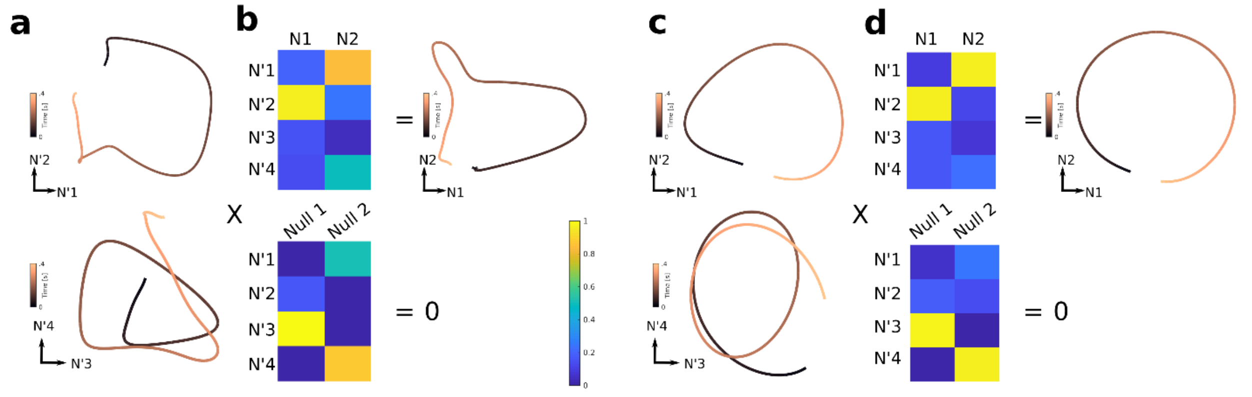


***Supplementary Fig. 6. Interpreting the null and potent spaces in presence of the Horseshoe effect*** *(**a**) First four PMd latent variables can be used to explain the activity in M1 and find potent and null spaces. The plots show the first and second (top), and third and fourth (bottom) leading PMd latent variables. (**b**) The colorplots shows the weight matrices that illustrate how the four leading latent variables of PMd are partitioned into a potent (top matrix) and a null (bottom matrix) subspace with respect to the two leading latent variables of M1 (plot on the right). The PMd potent space is predominantly limited to the two leading PMd latent variables. (**c-d**) Simulated data illustrate that the identical partition can be achieved without any relationship between the two datasets. Panels organization is identical to panels (a) and (b).*
